# Mathematical Modeling of Cancer Cell Invasion of Tissue: Development of an *in silico* Organotypic Assay

**DOI:** 10.1101/2025.05.12.653391

**Authors:** Vivi Andasari, Alf Gerisch, Andrew P. South, Mark A. J. Chaplain

## Abstract

**Background:** Organotypic assays are three-dimensional *in vitro* models widely used in cancer research to mimic the *in vivo* extracellular matrix (ECM) and to study cancer cell invasion by allowing investigation of critical interactions between tumor cells and their microenvironment. During invasion, cancer cells undergo genetic and epigenetic changes that disrupt cell-cell adhesion, enabling detachment from the primary tumor. Subsequently, invasive cells must (i) breach the basement membrane, a dense protein meshwork that restricts cell movement, and (ii) migrate through the surrounding ECM. This process relies on proteolytic enzyme secretion to degrade structural barriers, followed by cell-matrix adhesion-mediated migration.

**Methods:** We present an *in silico* model of cancer cell invasion in organotypic assays. The model is formulated as a system of partial differential equations capturing spatiotemporal dynamics, with nonlocal terms representing preferential adhesion-driven movement. Key variables in the model are cancer cells, proteolytic enzymes, and the ECM.

**Results:** Computational simulations demonstrate that modulation of cell-cell and cell-matrix adhesion parameters significantly influences tumor invasiveness. This result is consistent with experimental observations, demonstrating the model’s ability to accurately reflect *in vitro* behavior.

## 1 Introduction

Approximately 90% of cancer-related morbidity and mortality results from the dissemination of malignant tumor cells, a process known as metastasis. Metastasis occurs when cancer cells detach from the primary tumor, invade surrounding tissues, and colonize distant anatomical sites to form secondary tumors. This complex cascade depends on biochemical, biomechanical, and cellular interactions between tumor cells and the tumor stroma (the surrounding microenvironment).

Cancer invasion and metastasis involve a cascade of complex molecular processes connected through a series of adhesive interactions, proteolysis, and response to chemotactic stimuli. The major steps can be summarized as follows: (1) detachment of tumor cells from the primary tumor mass via downregulation of cell-cell adhesion molecules, such as cadherins and catenins; (2) local invasion through basement membrane and extracellular matrix (ECM) degradation by proteolytic enzymes, such as matrix metalloproteinases (MMPs) and urokinase-type plasminogen activator (uPA), and migration through the ECM, mediated by cell-matrix adhesion components such as integrins and fibronectin; (3) intravasation of the tumor cells into the blood vessels and lymphatic vessels, providing access to the circulatory system for travel to distant anatomical sites; (4) extravasation, where tumor cells move out of the blood vessels and into the surrounding tissue of new sites via adhesion to the endothelial cell lining at the capillary bed of the target organ site; and (5) colonization of target organs, culminating in secondary tumor growth [Guo and Giancotti, 2004, Brooks et al., 2010].

Our study focuses on local tissue invasion (steps 1 *−* 3), a critical phase in metastatic progression. Typically in carcinomas (epithelial-origin cancers), invasion involves stromal penetration and occupation by cancer cells that acquire genetic and epigenetic changes, leading to an aggressive phenotype. Aggressive and invasive (or malignant) cells often undergo epithelial-mesenchymal transition (EMT), where they lose their epithelial phenotypic properties and detach from epithelial sheets, transforming into mesenchymal cells. Increased cell motility through altered cell adhesion properties and the ability to release proteolytic enzymes are necessary for successful invasion. These properties are associated with decreased cell-cell adhesion, degradation of basement membranes and tissue, and enhanced cell-matrix adhesion for migration across the tissue that has been modified by proteolysis, as well as enhanced local growth of tumor cells. Continued invasion of the ECM occurs by repetition of these steps [Liotta, 1986].

### 1.1 The Role of Cell Adhesion in Cancer Progression

Cell adhesion is fundamental to developmental and pathological processes, including embryogenesis, wound healing, and cancer. Tumor cells exploit adhesion mechanisms to invade by losing cell-cell adhesion (to enable dissociation from primary tumors) and upregulating cell-matrix adhesion (to drive migration through degraded ECM). Dysregulated adhesion contributes to aggressiveness in multiple cancers, including colorectal [Kirkland and Ying, 2008, Paschos et al., 2009], breast [Kowalski et al., 2003, Sloan et al., 2006, Havaki et al., 2007], cervical [Maity et al., 2009], ovarian [Ahmed et al., 2005, Sawada et al., 2008], pancreatic [Sawai et al., 2006], lung [Lee et al., 2002, Zhu et al., 2007], carcinomas [von Schlippe et al., 2000, Brockbank et al., 2005, Heyder et al., 2005, Janes and Watt, 2006], and melanoma [Kuphal et al., 2005].

### 1.2 ECM Remodeling and Proteolytic Enzymes

The ECM acts as both a substratum on which cells move and a physical barrier that cells must overcome. One of the first steps in cancer invasion is ECM remodeling, a process that heavily involves the over-expression of proteolytic enzymes, such as urokinase-type plasminogen activator (uPA) and matrix metalloproteinases (MMPs). These enzymes, secreted by either the cancer cells themselves or the cells surrounding the tumor, degrade the physical obstacle by breaking down ECM proteins/components.

### 1.3 Organotypic Assays: Bridging *In Vitro* and *In Vivo* Models

Organotypic assays are three-dimensional *in vitro* systems developed to simulate/mimic *in vivo* ECM architecture by manipulating epithelial and stromal cells. In these assays, tumor cells are cultured atop a collagen I/Matrigel gel populated with human fibroblasts. The organotypic gel is then raised and maintained on a grid at the air/liquid medium interface, as shown in Fig. 1. Organotypic assays have been developed to mimic human tissue equivalents, particularly for cancer studies.

**Figure 1:**
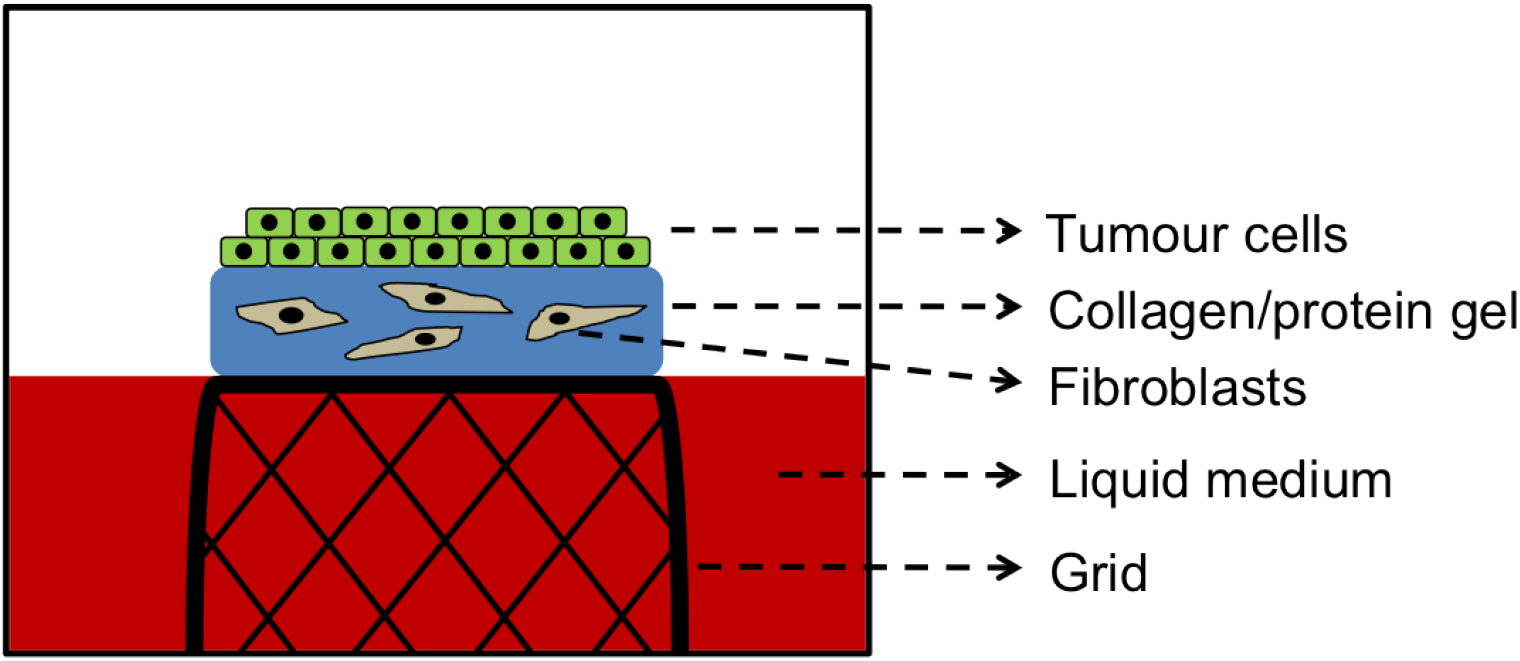
A schematic illustration of an organotypic assay used for tumor invasion studies.

One of the benefits of organotypic assays is their utility for basic studies of cell-cell and cell-matrix interactions. The similarity between invasion in organotypic assays and invasion observed in human tissues *in vivo* suggests that organotypic assays are physiologically relevant [Jenei et al., 2011]. Several *in vitro* studies using organotypic assays have produced results of invasion and differentiation similar to those observed in skin cancer [Akgül et al., 2005, Nyström et al., 2005, Martins et al., 2009, Jenei et al., 2011, Moutasim et al., 2011], breast cancer [Holliday et al., 2009], oral cancer [Brusevold et al., 2010a,b], pancreatic cancer [Froeling et al., 2009, 2010], and prostate cancer [Papini et al., 2007].

### 1.4 Objectives and Model Overview

In this paper, we develop an *in silico* model of cancer cell invasion that integrates cell-cell/cell-matrix adhesion, proteolytic ECM degradation, and cell motility, based on *in vitro* models using organotypic assays. Our mathematical model consists of a system of three partial differential equations representing cancer cell density, ECM density, and matrix proteolytic enzyme concentration. The movement of the cancer cell population is governed by random motion, adhesion-mediated migration involving cell-cell and cell-matrix adhesion, and chemotaxis due to the gradient of matrix-degrading enzyme concentration.

In the **Results and Discussion** section, we present computational simulations that exhibit qualitative comparisons with experimental data. In the **Conclusions** section, we compare the computational simulation results of our *in silico* model with an *in vitro* model presented in Martins et al. [Martins et al., 2009]. We also briefly discuss a wider range of patterns that emerge by varying parameter values of the mathematical model.

## 2 Methods

The organotypic assay, originally developed by Fusenig and colleagues Fusenig et al. [1983], is a 3D cell-migration assay widely used to study the invasive potential of various cancer types. This assay consists of a 3D collagen gel embedded with fibroblasts, serving as a simulated ECM. Cancer cells are seeded atop the gel and allowed to migrate vertically into the gel to varying degrees of penetration. The invasion depth is quantified via an “Invasion Index” Nyström et al. [2005], a metric that combines the average invasion depth, the number of tumor islands Jenei et al. [2011], and the area of invading tumor islands.

Cell adhesion is a critical driver of invasion. Continuum models have been developed to incorporate cell adhesion, such as models that incorporated cell adhesion via a boundary condition as a surface tension-like force Greenspan [1976], Bell et al. [1984], Evans [1985a,b], Bell and Torney [1985], Hammer and Lauffenburger [1987], DiMilla et al. [1991], Xiao and Truskey [1996], Byrne and Chaplain [1996], Byrne [1997], Byrne and Chaplain [1997], Palecek et al. [1997], Cristini et al. [2005], Zheng et al. [2005], Macklin and Lowengrub [2007], Cristini et al. [2009], Frieboes et al. [2010], or models where cell adhesion was modeled in implicit terms Perumpanani et al. [1996]. The first continuum model using PDEs for adhesion-driven cell migration was introduced by Armstrong et al. Armstrong et al. [2006], with key features: (i) a nonlocal adhesion term that accounts for forces within a “sensing radius,” (ii) extensions to cancer invasion models incorporating cell-matrix adhesion Gerisch and Chaplain [2008], Painter et al. [2010], and (iii) theoretical foundations with boundedness of solutions Sherratt et al. [2009].

### 2.1 Model Formulation

In our *in silico* organotypic assay model, we consider a cancer cell invasion model with three time- and space-dependent variables:

- *c*(*t*, **x**): cancer cell density,
- *v*(*t*, **x**): density of stroma or ECM,
- *m*(*t*, **x**): concentration of matrix proteolytic enzyme (*e*.*g*., MMP-2).

The vector of these three variables in a compact notation is defined as **w**(*t*, **x**) := (*c*(*t*, **x**), *v*(*t*, **x**), *m*(*t*, **x**)), where the temporal variable *t ∈* [0, *T*] and the spatial variable 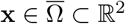 (2D domain).

### 2.2 Governing Equations

Since cancer cell invasion depends on the interplay between adhesive and proteolytic mechanisms, our model accounts for cell adhesion and chemotaxis in the movement of cancer cells within an organotypic assay.

- **Cancer cell dynamics:** Cancer cell migration is governed by random motion (with cell random motility coefficient *D*_1_, adhesive movement, and chemotaxis due to matrix proteolytic enzyme (with chemotactic sensitivity coefficient *ξ*_*m*_. We assume a cancer cell proliferation term in the form of a logistic growth law with proliferation parameter *φ*_1_.

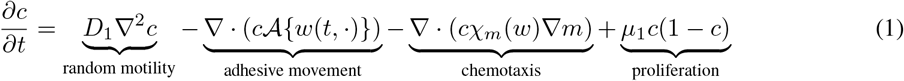
- **ECM degradation:** Changes in the collagen distribution of the organotypic assay are assumed to result from degradation upon contact with the matrix proteolytic enzyme at a degradation rate *β*. Since the matrix does not diffuse, there is no diffusion term in this equation.

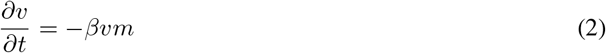
- **Protease dynamics:** The spatio-temporal evolution of the matrix proteolytic enzyme concentration is assumed to occur through diffusion with diffusion coefficient *D*_3_, production by cancer cells at rate *γ*, and natural degradation at rate *σ*.

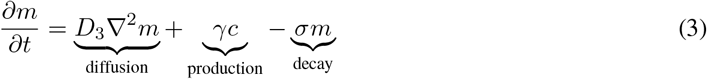

### 2.3 Adhesion Velocity

The advection term 𝒜{**w**(*t,·*,)} in Eq. (9a) represents the “nonlocal” adhesion velocity. We adapt the form for 𝒜{**w**(*t,·*,)} introduced by Armstrong et al. Armstrong et al. [2006]. This form has been studied and applied in various cancer models Gerisch and Chaplain [2008], Sherratt et al. [2009], Painter et al. [2010]. Following Gerisch and Chaplain Gerisch and Chaplain [2008], for 1-D spatial domains, the form of 𝒜{**w**(*t,·*,)} is given by

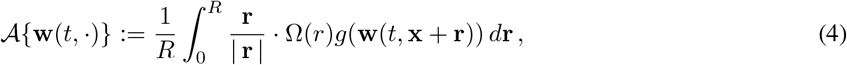

where the forces generated through adhesive binding affect the points in the right 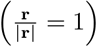or left 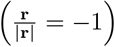 direction of the Cartesian coordinate axis. For 2-D spatial domains, the adhesion velocity is given by

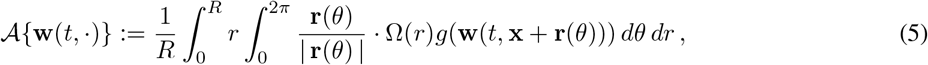

where the forces are sensed by points within a ball of radius *R*. The position vector (or the outer normal vector) **r** in polar coordinates is **r**(*θ*) = *r*(cos *θ*, sin *θ*)^*t*^*op* and its magnitude is |**r**| = *r*.

The function Ω(*r*) describes the strength of the adhesion velocity 𝒜{w(*t,·*,)} as influenced by points within the sensing region at **x**, depending on their distance **r** from **x**. Several forms of Ω(*r*) have been discussed, and the reader is referred to Armstrong et al. [2006], Gerisch and Chaplain [2008], Painter et al. [2010] for a comprehensive description. For simplicity, in this paper, we consider the following constant form for 2D spatial domains,

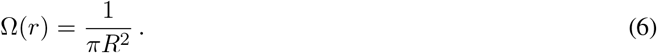

This form implies that the strength of the adhesion velocity is uniform in all directions, irrespective of the distance.

### 2.4 Volume Filling and Chemotaxis

Since adhesion involves forces between cancer cells (cell-cell adhesion) and between cancer cells and stroma (cell-matrix adhesion), we define *g*(**w**(*t*, **x** + **r**)) as a function of *c* and *v*. For convenience, we write <span class=“math-inline”>g*c, v*</span>. We consider a form for *g*(*c, v*) that incorporates a spatial restriction, where cells aggregate and move only if there is space locally available

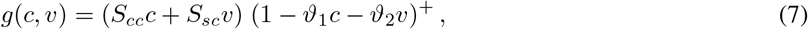

as discussed in Gerisch and Chaplain [2008]. The factor (1 *−ϑ*_1_*c −ϑ*_2_*v*)^+^ := max {0, (1 *−ϑ*_1_*c −ϑ*_2_*v*)} ensures that the force generated is limited by the densities of cells and matrix. We assume that in one unit volume of physical space, cells occupy a fraction *ϑ*_1_ of the space, and the stroma occupies another fraction *ϑ*_2_.

In another cancer invasion model analyzed in our previous work Andasari et al. [2011], we considered a constant form for the chemotactic sensitivity coefficient, *χ*_*m*_(**w**) = *χ*_*m*_. This model has been intensively studied in the mathematical biology and applied PDE literature. The constant form of *χ*_*m*_ means that the gradient of *∇u* is multiplied by a linear, rather than a nonlinear, function. One drawback of this choice is that it may lead to finite-time blow-up or “overcrowding” scenarios, where the solution becomes unbounded.

The linear form can also pose difficulties for numerical simulations. Therefore, we opt for a form that includes assumptions regarding spatial restrictions, such as limiting the movement of cells at relatively high densities. One way to achieve this is by switching off the chemotactic response when the cell density becomes high, preventing cells from moving into dense regions. This mechanism, also called “overcrowding prevention” or “volume filling,” has been studied by Hillen and Painter Hillen and Painter [2001], Painter and Hillen [2002] and references therein. We use the same form as the function *g*(*c, v*) in Eq. 7, and the chemotactic sensitivity coefficient *χ*_*m*_(**w**) in a non-dimensional form is given by

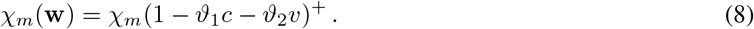

### 2.5 Non-Dimensionalization

The system of equations in Eqs. (9a)-(9c) can be non-dimensionalized using the following scales:

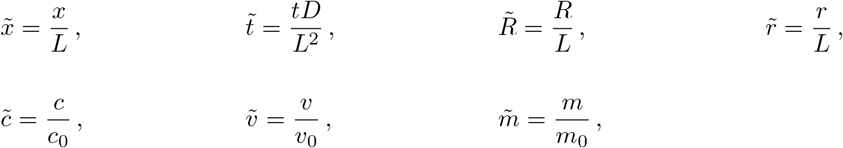

where, as in Andasari et al. [2011], *L* := 0.1 cm is a characteristic length scale, representing the maximum distance of the cancer cells at the early stage of invasion, *D* := 10^*−*6^ cm^2^s^*−*1^ is a representative chemical diffusion coefficient, and *τ* = *L*^2^*/D* = 10^4^s. The dependent variables *c, v*, and *m* are rescaled with *c*_0_ = 6.7 *×* 10^7^ cells cm^*−*3^, *v*_0_ = 1 nM, and *m*_0_ = 0.1 nM, respectively. The parameters 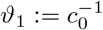 and 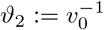 represent the space fraction per unit cancer cell density (1.5 *×*10^*−*8^ cm^2^ cell^*−*1^) and the space fraction per unit density of matrix (1 nM^*−*1^), respectively Gerisch and Chaplain [2008]. The typical radius of a cell without protrusion is between 5 *−* 50 *μ*m. This radius is distinct from the sensing radius *R*. To reach its surroundings, a cell extends its body (or cellular membranes) to form protrusions. The protrusion or the maximum body extension of a cell can reach 2 to 10 times its radius. This means that the scaling factor is *η ∈* [0.1, 0.5] and hence *R* is approximately between 10 *−* 100 *μ*m Xue et al. [2014], and the dimensionless 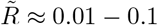

We derive the dimensionless system, with tildes dropped for clarity, as

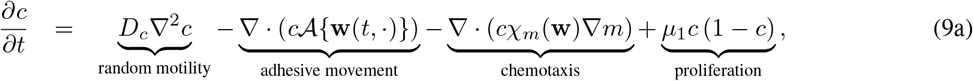

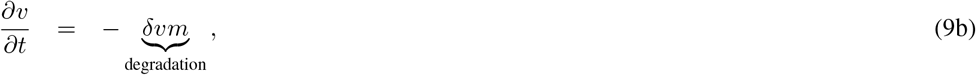

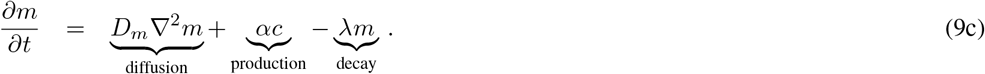

### 2.6 Numerical Implementation

The system of equations (9) is closed with appropriate initial and boundary conditions. The *in vitro* organotypic models, such as those in Martins et al. [2009], are 3D. However, the analysis is carried out on 2D cross-sections (see Fig. 2). Therefore, we adopt a 2D model, and our computational simulation results can be directly compared with the experimental analysis.

**Figure 2:**
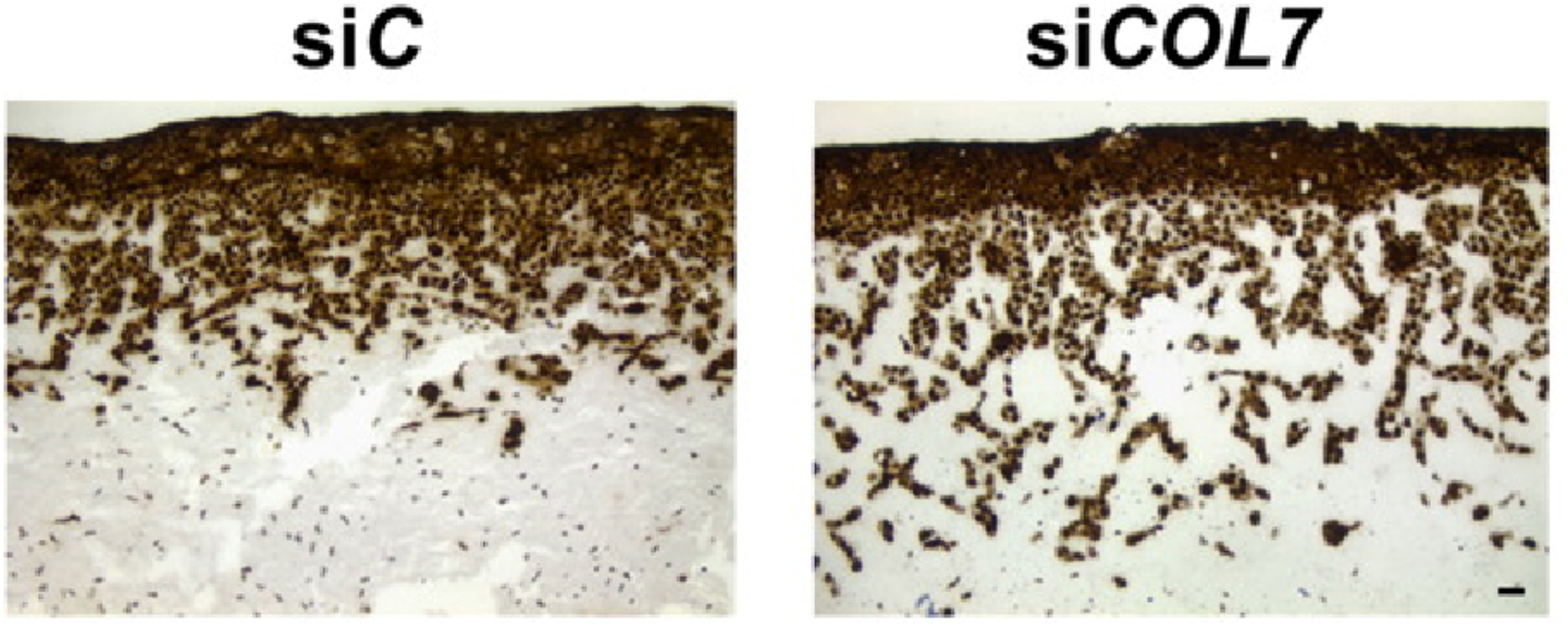
Figures showing organotypic cultures of siCOL7 and siC MET1 cells on collagen:matrigel gels with incorporated fibroblasts immunostained with a pan-cytokeratin antibody and visualised with DAB from experimental results by Martins et al. Martins et al. [2009]. siCOL7 cells displayed increased invasion into the gel compared with siC cells. Reproduced with permission from the publisher of Martins et al. Martins et al. [2009].

For the initial condition, following the organotypic assay setup shown in Fig. 1, we assume that some cancer cells are initially present in the domain, seeded atop it as a strip of uniform initial density. This initial strip of cancer cells occupies approximately 12.5% of the domain with a density of 1, while the matrix concentration uniformly occupies the remaining 87.5% of the domain, also with an initial concentration of 1. The initial concentration of matrix proteolytic enzyme is half of the initial concentration of the cancer cells.

For 2D simulations, we use the spatial domain *D*_2_ := (0, *M*_*x*_) × (0, *M*_*y*_) ⊂ ℝ with positive parameters *M*_*x*_ = 3 (the length of the domain in the *x*-direction) and *M*_*y*_ = 2 (the length of the domain in the *y*-direction). We impose zero-flux boundary conditions on all sides of the rectangular domain. The schematic for the setup of initial and boundary conditions for the 2D domain of computational simulations is shown in Fig. 4.

In the experiments by Martins et al. Martins et al. [2009], squamous cell carcinoma (SCC) cells with depleted type VII collagen (ColVII) exhibit enhanced SCC migration and invasion (see Fig. 2). ColVII is a key component of anchoring fibrils, responsible for the attachment between the external epithelia and the underlying stroma. Fig. 3 shows that SCC cells lacking ColVII display: (i) decreased expression of membranous E-cadherins, (ii) increased expression of *α*_*v*_*β*_3_ integrins in invading ColVII knockdown epithelial cells, and (iii) an increase in vimentin-positive cells at the invasion front. Vimentin intermediate filaments are thought to be associated with *α*_*v*_*β*_3_ integrin-rich endothelial focal contacts. E-cadherins are the primary adhesion molecules mediating cell-cell adhesion, whereas integrins are associated with cell-matrix adhesion. To investigate the influence of adhesive movement and chemotaxis in our model, particularly the effects of cell adhesion parameters, we reduce the influence of random motion by using small diffusion coefficients. We hypothesize that an invasive situation is best represented by nonlocal parameter values where the cell-matrix adhesion coefficient *S*_*cv*_ is higher than the cell-cell adhesion coefficient *S*_*cc*_. Biologically, this suggests that cells increase their attachment to the matrix/stroma to facilitate movement across it and reduce their attachment to other cells, allowing them to detach and migrate. Conversely, to model scenarios with less invasion, the cell-cell adhesion coefficient *S*_*cc*_ is set to be higher than the cell-matrix adhesion coefficient *S*_*cv*_.

**Figure 3:**
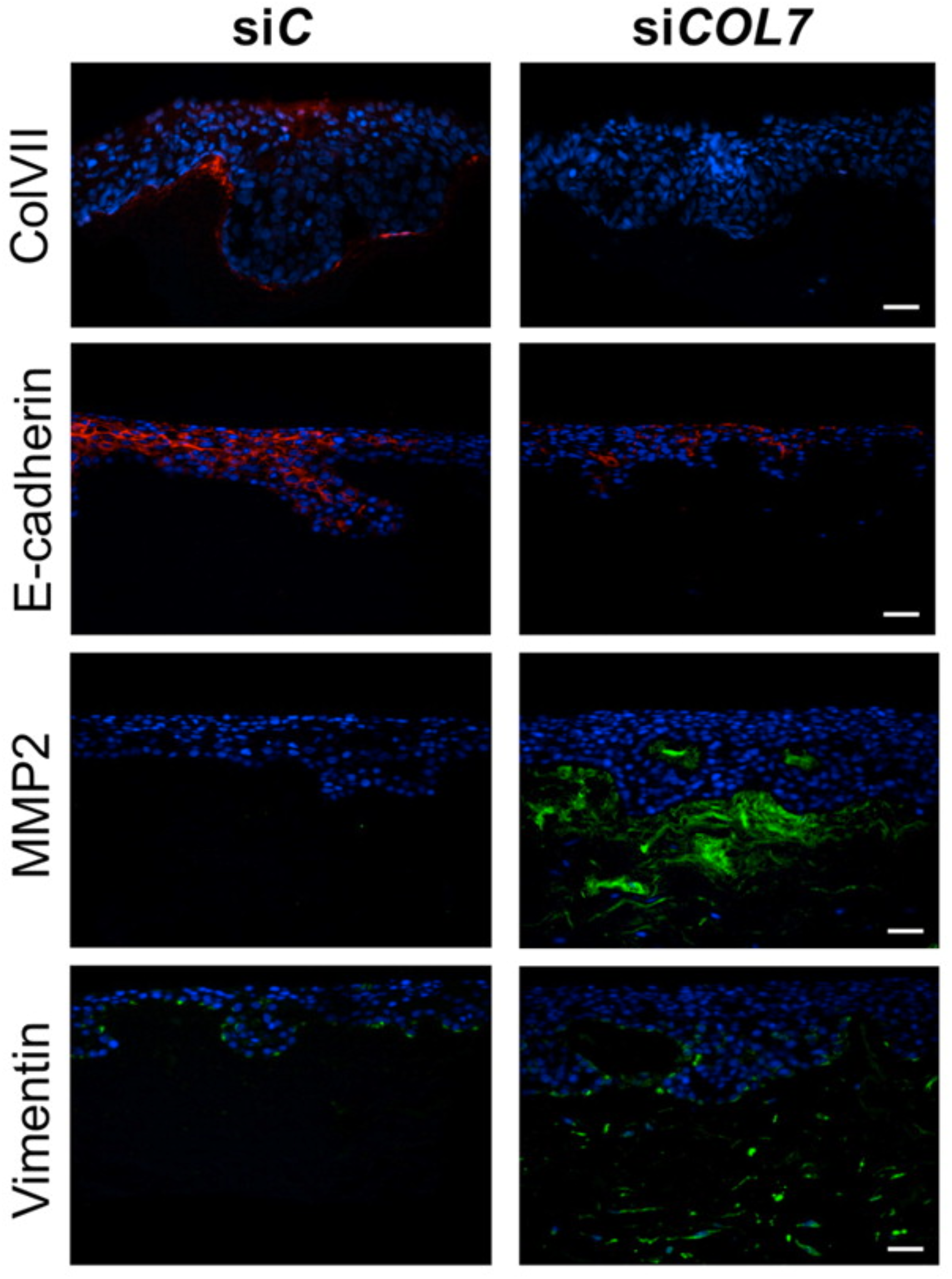
Figures showing immunofluorescence staining of sections of siC and siCOL7 MET1 organotypic cultures on de-epidermalised dermis (DED) with antibodies to ColVII (red) and the EMT markers, E-cadherin (red), MMP2 (green), and vimentin (green) from the experimental results by Martins et al. Martins et al. [2009]. The results show a decreased expression of membranous E-cadherin and increased expression of MMP2 and vimentin-positive cells in the papillary dermis in siCOL7 sections. Reproduced with permission from the publisher of Martins et al. [2009].

**Figure 4:**
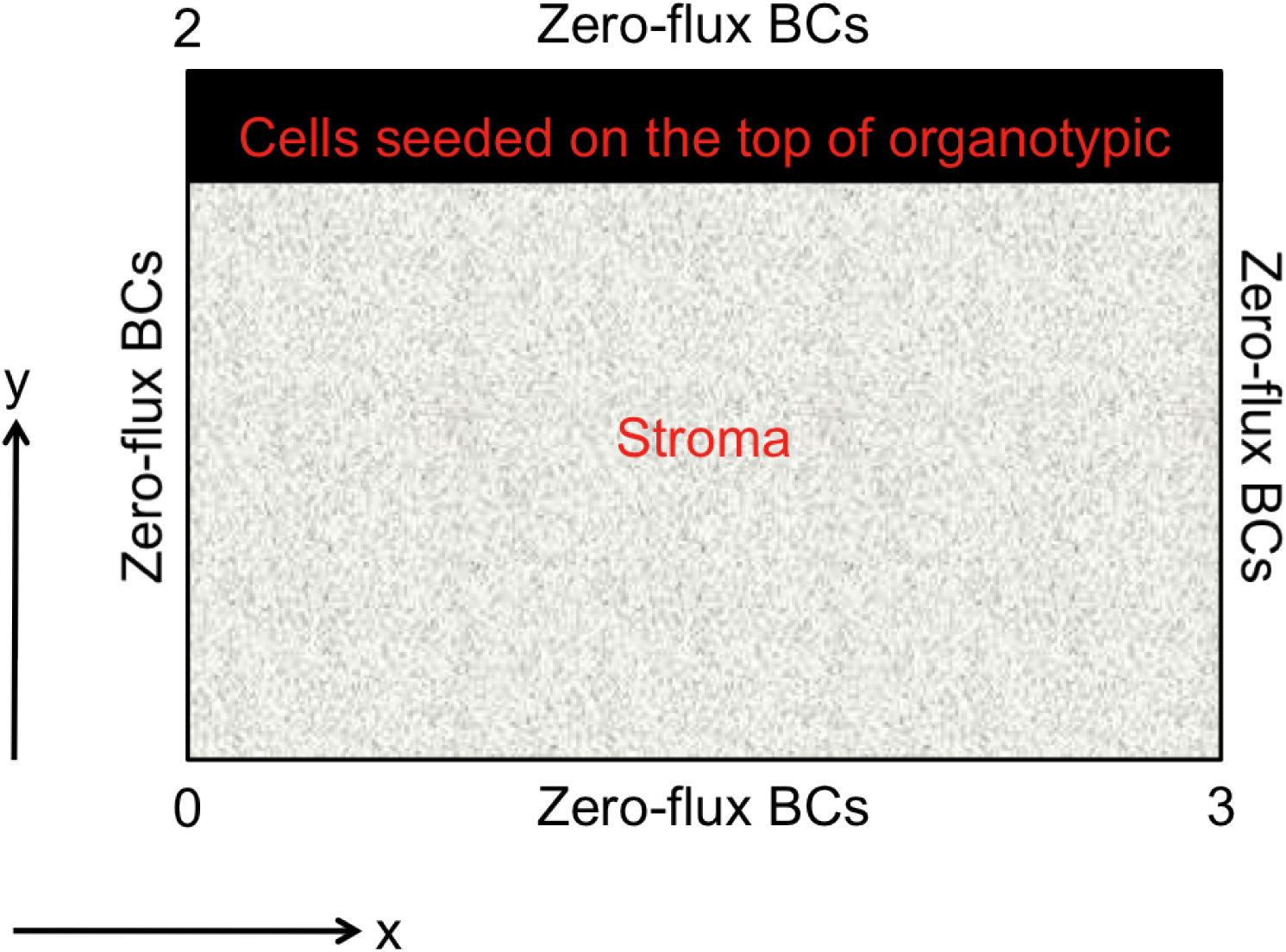
Schematic diagram showing the initial and boundary conditions used for 2D computational simulations on a rectangular domain *D* = (0, 3) *×* (0, 2). Zero-flux boundary conditions are applied on all boundaries.

Sun et al. Sun et al. [2004] measured a cell random motility coefficient of 0.07 *μ*m^2^*/*min by observing human fibroblast migration in a 3D collagen gel. Using a reference diffusion coefficient *D* := 10^*−*6^ cm^2^s, this measurement yields a dimensionless cancer cell random motility coefficient on the order of *D*_*c*_ *∼* 10^*−*5^. This value should be sufficient for our computational simulations, as our primary aim is to focus on the role of adhesion in driving cancer cell movement or invasion. By employing smaller diffusion coefficients, we emphasize adhesion-driven movement and aim to induce heterogeneity in the solutions, as the volume filling mechanism incorporated in the adhesive and chemotaxis movement terms might otherwise stabilize the homogeneous steady state, resulting in traveling wave-like solutions. Our goal in this paper is to demonstrate the diverse patterns that emerge in the solutions when we vary cell-cell and cell-matrix adhesion parameters. Furthermore, it has been observed that varying cell-cell and cell-matrix adhesion can affect the speed of cell dispersion or invasion. This observation supports the hypothesis by Fidler Fidler [1978] that the adhesive properties of cancer cells can influence the patterns of spread and growth. The specific patterns that evolve are governed by the full nonlinear system; therefore, a nonlinear stability analysis could potentially provide further insight into these effects, although we do not perform such an analysis in this paper.

### 2.7 Parameterization

In this work, we consider a default set of model parameters, referred to as *parameter set 𝒜*, given by

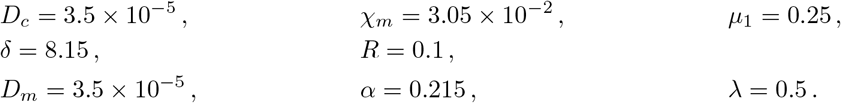

Most of these parameter values were adopted from our previous work Andasari et al. [2011]. In this study, we specifically vary the adhesion parameters (*S*_*cc*_, *S*_*cv*_) to investigate their effects, as mentioned earlier.

### 2.8 Model Extensions

The extended system of equations, incorporating an additional partial differential equation for fibroblast cells, can be written as

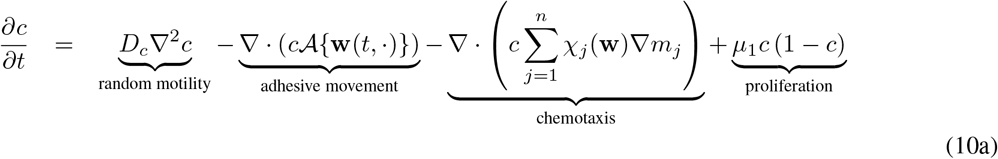

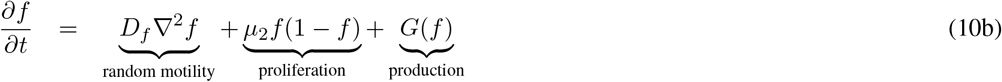

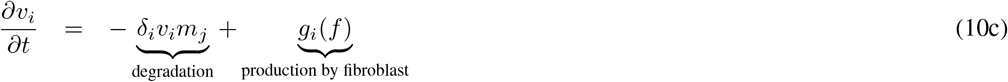

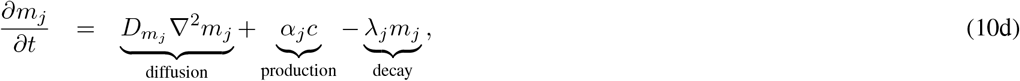

where *i* = 1, 2, …, *p* and *j* = 1, 2, …, *n* depend on the number of matrix components *v*_*i*_ and matrix proteolytic enzymes *m*_*j*_ included in the model. In this extended model, the function *g*(**w**) in the adhesive movement term *𝒜{w(t,·,)}* incorporates an additional adhesive component due to the fibroblasts,

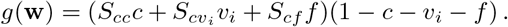

Finally, we note that future development of this modeling approach (an *in silico* organotypic assay) should involve attempts to estimate the model parameters from a single experimental setup and to quantify our adhesion parameters, specifically *S*_*cc*_ and *S*_*cv*_. Achieving this would enable the development of a quantitative modeling approach that would be highly valuable for experimentalists.

## 3 Results and Discussion

We solved the system (9) numerically using the initial and boundary conditions described in the **Methods** section. The numerical simulations were performed using the same numerical technique as in Andasari et al. [2011], with an additional evaluation for the integral terms. The numerical scheme employs the method of lines, where spatial discretization utilizes a second-order finite volume approach incorporating flux-limiting for accurate discretization of the chemotaxis and nonlocal terms. Specifically for the nonlocal terms, spatial discretization is achieved using the Fast Fourier Transform (FFT) technique. Comprehensive details of the numerical scheme are provided in Gerisch [2010], Gerisch and Chaplain [2006, 2008]. Unless stated otherwise, the parameters used are from parameter set *A*.

### 3.1 Spatial Domains

The simulations for a 2D spatial domain were conducted on a rectangular domain *D* := (0, *M*_*x*_) × (0, *M*_*y*_) ⊂ ℝ^2^, where *M*_*x*_ = 3 and *M*_*y*_ = 2 in the *x* and *y* directions, respectively, for all simulations unless otherwise specified.

Each unit dimension was discretized into 200 grid cells, resulting in Δ*x* = 0.015 and Δ*y* = 0.01. We previously simulated the system using first 100 grid cells (Δ*x* = 0.03, Δ*y* = 0.02) and then 150 grid cells (Δ*x* = 0.02, Δ*y* = 0.0133). The simulation results were qualitatively consistent across these grid resolutions. However, upon refining the grid, minor quantitative differences, such as the precise depth of cancer cell penetration, were observed. We found that further refinement beyond 200 grid cells yielded simulation results indistinguishable from those with 200 grid points, and therefore, we adopted this grid size for all subsequent simulations. We ran the simulations until the solutions approached the bottom boundary (*y* = 0). Initially, at *t* = 0, we assumed a uniform cell density occupying 1*/*8 of the domain’s area, with the extracellular matrix occupying the remaining area, as depicted in Fig. 4.

### 3.2 Adhesion-Driven Invasion Patterns

In Fig. 5, we present solutions for cancer cell density at dimensionless time *t* = 100, using parameter set 𝒜. We varied the cell-cell (*S*_*cc*_) and cell-matrix adhesion (*S*_*cv*_) parameters to model an “invasive scenario” and a less invasive scenario, enabling a comparison of our *in silico* results with the *in vitro* results reported by Martins et al. Martins et al. [2009]. For the less invasive scenario, we set *S*_*cv*_ = 0.01 <*S*_*cc*_ = 0.1 (cell-matrix adhesion less than cell-cell adhesion), computationally replicating the left panel of Fig. 2 (siC). Our corresponding *in silico* simulation result is shown in the left-hand figure of Fig. 5. Consistent with the *in vitro* experimental observation, we observe relatively slow movement of cancer cell density, attributed to the cells maintaining a stronger attachment to neighboring cells and a weaker attachment to the stroma/matrix.

**Figure 5:**
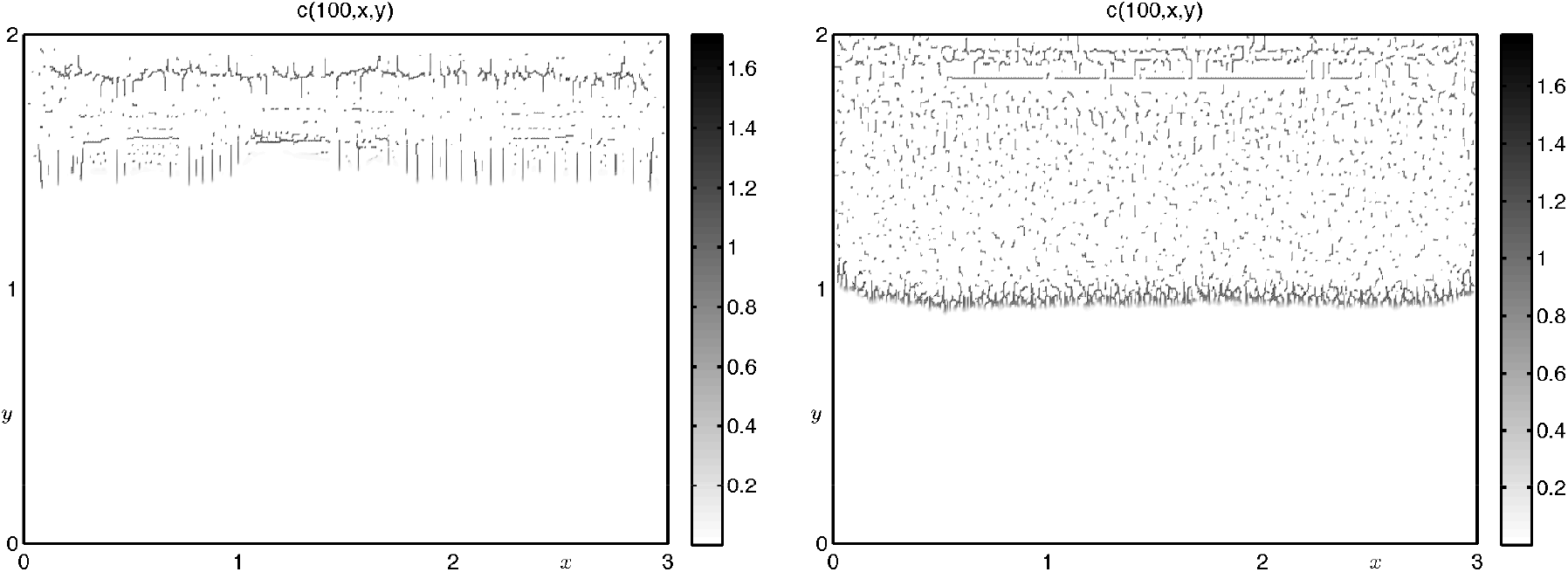
Plots showing the distribution of the cancer cell density *c*(*t, x, y*) using parameter set 𝒜 with *S*_*cv*_ = 0.01, *S*_*cc*_ = 0.1 (left) for the less invasive scenario and *S*_*cv*_ = 0.1, *S*_*cc*_ = 0.01 (right) for the invasive scenario taken at dimensionless time *t* = 100.

To model the more invasive scenario, as depicted in the right panel of Fig. 2 (siCOL7), we set *S*_*cv*_ = 0.1 *> S*_*cc*_ = 0.01, representing stronger cell-matrix adhesion. The corresponding *in silico* result is shown in the right-hand figure of Fig. 5. We observe that the cancer cells invade more deeply into the domain compared to the less invasive scenario, which aligns with the experimental findings of Martins et al. Martins et al. [2009]. Our *in silico* results also exhibit qualitative similarities to experimental results obtained in other organotypic assays by Nystrom et al. Nyström et al. [2005] and Jenei et al. Jenei et al. [2011].

Quantification of the invasion depth in our *in silico* results reveals that cells in the invasive scenario simulation penetrated approximately twice as deep into the stroma as cells in the less invasive scenario. This quantitative difference is comparable to the measurements of invasion depth reported in Martins et al. [2009]. Notably, this difference in invasion depth was achieved in our model by altering only the cell-cell and cell-matrix adhesion parameters, highlighting the significant role of these parameters in regulating cancer cell invasiveness.

Other intriguing invasive patterns emerge when the cell-cell and cell-matrix adhesion parameters are varied. When both parameters are reduced to *S*_*cc*_ = *S*_*cv*_ = 0.01, we observe less aggregation and reduced invasion of cell density patterns, as illustrated in the right panel of Fig. 6. Conversely, increasing both cell-cell and cell-matrix adhesion to an equally strong level, *S*_*cc*_ = *S*_*cv*_ = 0.1, results in a distinctly different pattern of cell density, as shown in the left panel of Fig. 6. The stronger overall adhesion leads to faster and deeper invasion of the cancer cells, suggesting that the absolute strength of adhesion can also modulate the dynamics of invasion.

**Figure 6:**
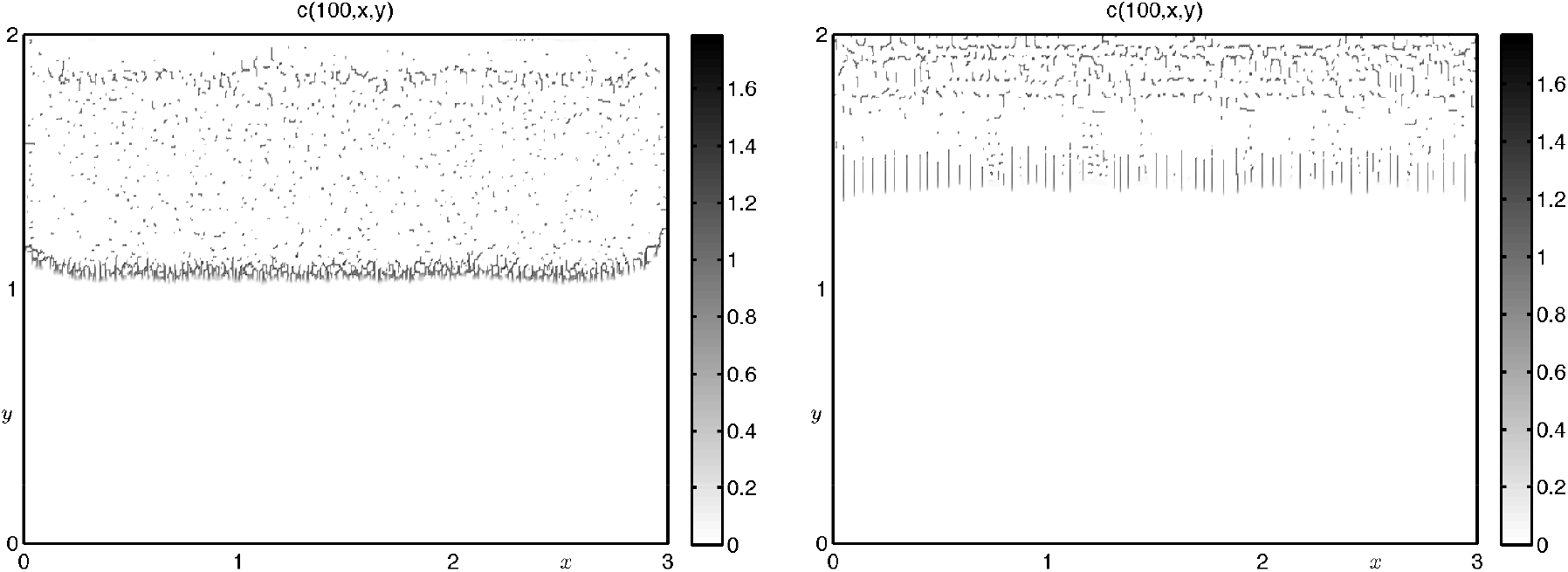
Plots showing the distribution of the cancer cell density *c*(*t, x, y*) using parameter set 𝒜 with *S*_*cv*_ = 0.1, *S*_*cc*_ = 0.1 (left figure) to represent equally strong cell-cell and cell-matrix adhesion and *S*_*cv*_ = 0.01, *S*_*cc*_ = 0.01 (right figure) to represent equally weak cell-cell and cell-matrix adhesion, taken at dimensionless time *t* = 100.

### 3.3 Chemotaxis Modulation

We next examine the effects of varying the chemotactic parameter *χ*_*m*_, while keeping all other parameters as in parameter set 𝒜. The simulation results are presented in Fig. 7, where *χ*_*m*_ = 0.01 represents reduced chemotaxis (left figure) and *χ*_*m*_ = 0.05 represents increased chemotaxis (right figure). In both cases, cell-matrix adhesion was stronger than cell-cell adhesion (*S*_*cv*_ = 0.1 and *S*_*cc*_ = 0.01). The results suggest that modulating chemotactic activity can significantly influence the development of a more metastatic phenotype or invasive behavior. These findings are consistent with the experimental results of Martins et al. Martins et al. [2009], where SCC invasion and epithelial-mesenchymal transition (EMT) are associated with the overexpression of matrix proteolytic enzymes, such as MMP-2. Specifically, the experimental results shown in the third panel from the top of Fig. 3 demonstrate an increased expression of MMP-2.

**Figure 7:**
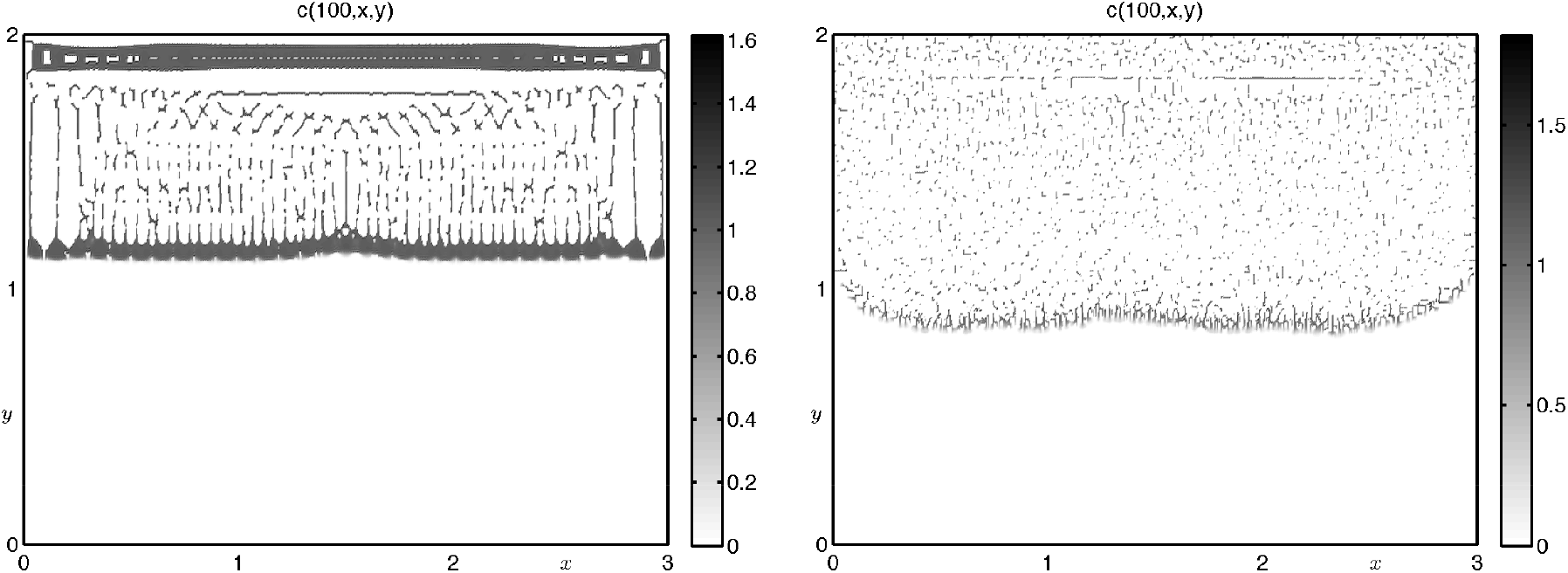
Plots showing the distribution of the cancer cell density *c*(*t, x, y*) using parameter set 𝒜 with weaker chemotactic strength (*χ*_*m*_ = 0.01, left figure) and stronger chemotactic strength (*χ*_*m*_ = 0.05, right figure), both with the same adhesion parameters, *S*_*cv*_ = 0.1, *S*_*cc*_ = 0.01, taken at dimensionless time *t* = 100.

### *3*.*4 In Silico* Scratch Assays

Another experimental method used to study and quantify cell migration *in vitro* is the scratch assay (also known as the wound healing assay). These assays are performed in a 2D monolayer by creating a “scratch” or “wound” in the cell monolayer and then monitoring cell migration across the created gap. This technique is considered particularly suitable for studying the effects of cell adhesion, but less suitable for studying chemotaxis Moutasim et al. [2011]. Fig. 8 shows an *in vitro* scratch assay experiment by Martins et al. Martins et al. [2009], where SCC cells lacking ColVII (siCOL7) and control cells with ColVII (siC) were plated on a cell monolayer. After 36 hours, siCOL7 cells invaded the cell monolayer and closed the “scratch,” while the siC cells exhibited less invasive/migratory activity.

**Figure 8:**
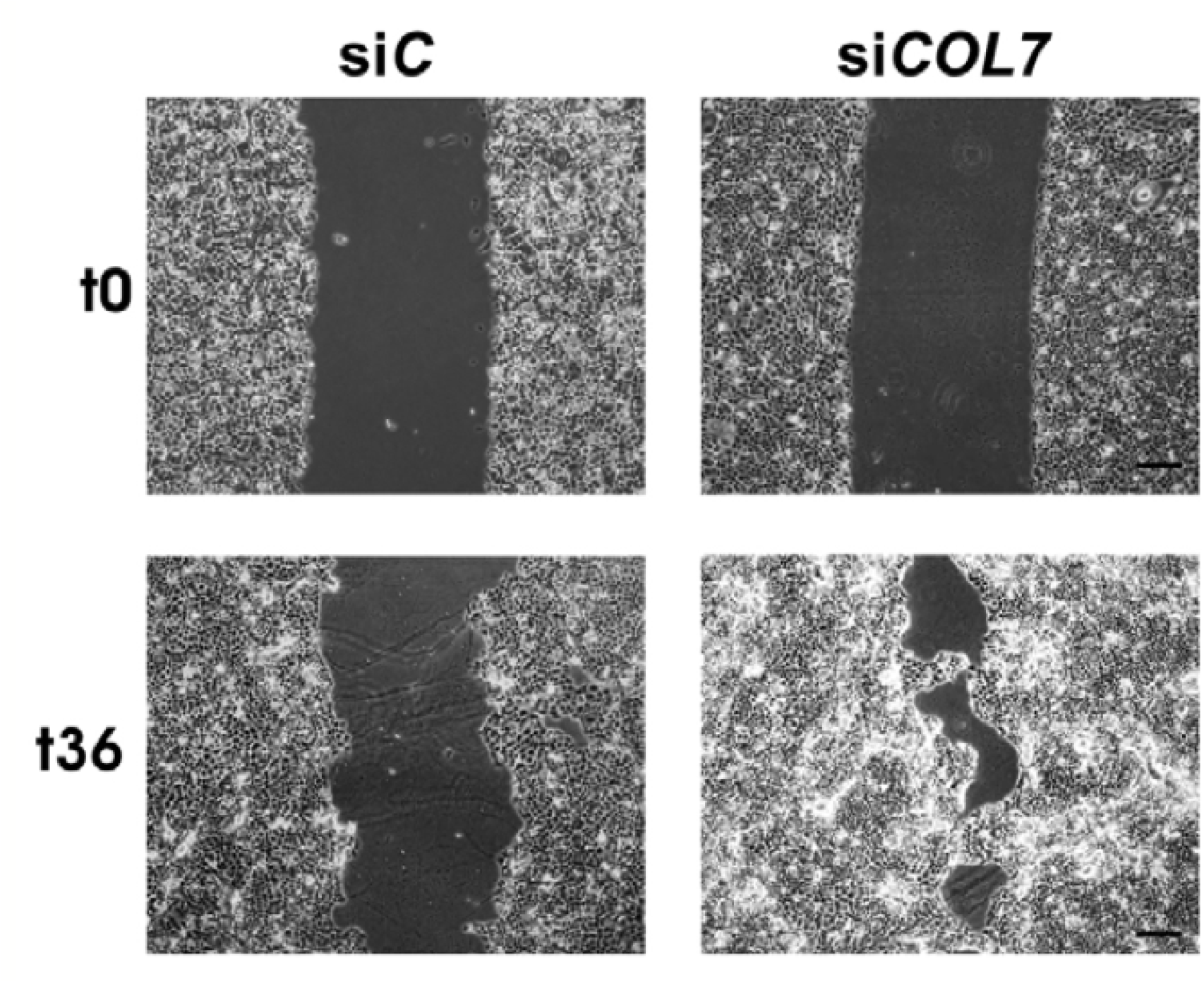
Experimental results of Martins et al. [2009] showing an *in vitro* scratch assay using siC and siCOL7 MET1 cells, taken before migration (t0, top figures) and 36 hours after migration (t36, bottom figures). The figures show that the siCOL7 cells migrate/invade more rapidly than the siC cells. Reproduced with permission from the publisher of Martins et al. [2009].

Given that the *in vitro* scratch assay experiment in Fig. 8 is 2D, we can readily adapt our basic invasion model to create an *in silico* scratch assay and model cell invasion/migration in this setting. We modify our initial conditions to mimic the scratch assay setup: the initial cancer cell density is set to one and occupies 2*/*3 of the domain, divided into two strips, with 1*/*3 placed on the right side and 1*/*3 on the left side. The initial stroma density is also set to one and occupies the remaining 1*/*3 of the domain in the center. The initial concentration profile of proteolytic enzyme is half that of the cell density. The initial conditions are illustrated in Fig. 9.

**Figure 9:**
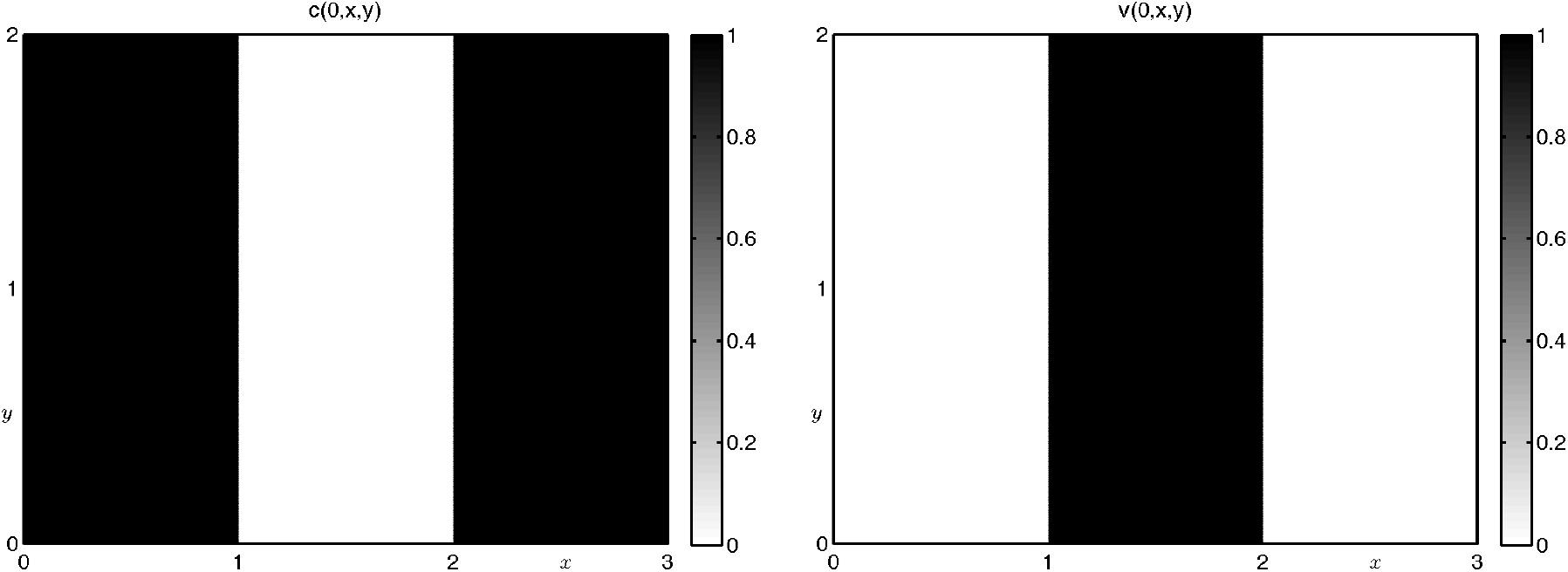
Plots showing the initial conditions of cancer cell density (left figure) and stroma (right figure) for *in silico* scratch assay simulations.

The computational results at dimensionless time *t* = 55 are shown in Fig. 10. The left panel depicts stronger cell-cell adhesion (*S*_*cv*_ = 0.01 and *S*_*cc*_ = 0.1), and it is evident that the cells exhibit very slow movement into the central gap (see Fig. 8). In contrast, the right-hand panel, representing stronger cell-matrix adhesion (*S*_*cv*_ = 0.1 and *S*_*cc*_ = 0.01), shows that the cancer cells invade/migrate rapidly into the gap and almost connect in the middle of the domain (Fig. 8). Again, these distinct outcomes are achieved solely by varying the adhesion parameters, emphasizing their critical role in dictating the migratory behavior of cancer cells.

**Figure 10:**
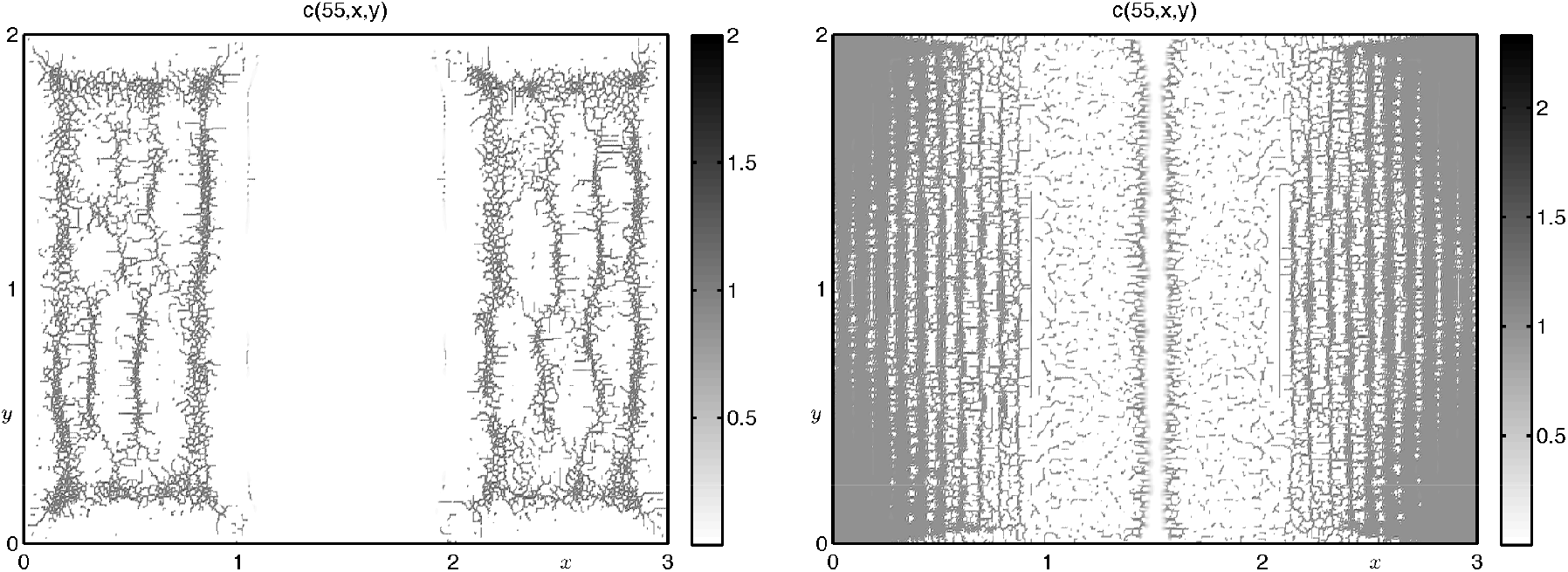
Plots showing the cancer cell density *c*(*t, x, y*) migrating in an *in silico* scratch assay under two different adhesion scenarios. The left figure shows the extent of cell migration/penetration using parameter set 𝒜 with *S*_*cv*_ = 0.01, *S*_*cc*_ = 0.1 for a less invasive scenario. The right figure shows the extent of cell migration/penetration using parameter set 𝒜 with *S*_*cv*_ = 0.1, *S*_*cc*_ = 0.01 for a more invasive scenario. Both plots are taken at dimensionless time *t* = 55. The figures show that changing only the adhesion parameters in our model can give rise to completely different invasion results in line with experimental results.

### 3.5 Pattern Formation at Higher Motility

The model also exhibits a wide range of other “cellular patterns” depending on the choice of parameters. To explore the effects of increased cell random motility and matrix-degrading enzyme diffusion, we performed computational simulations using higher coefficients: *D*_*c*_ = 3.5 × 10^*−*4^ and *D*_*m*_ = 3.5× 10^*−*4^, which are ten times greater than the corresponding parameters in set 𝒜. These simulations were conducted on a square spatial domain *D*_2_ := (0, 3) × (0, 3) with the same initial and boundary conditions as described in the **Methods** section. The results, shown in Fig. 11, reveal a variety of intriguing spatiotemporal patterns obtained by varying the chemotactic coefficient and cell adhesion parameters. The top left panel (a) illustrates a “vertical stripe” pattern, generated with *χ*_*m*_ = 0.04 and *S*_*cc*_ = *S*_*cv*_ = 0.1. The top right panel (b) of Fig. 11 displays a “horizontal stripe” pattern, obtained with *χ*_*m*_ = 0.04 and *S*_*cc*_ = *S*_*cv*_ = 0.01, while the lower figure (c) shows a “mixed horizontal and vertical stripe” pattern resulting from *χ*_*m*_ = 0.0305 and *S*_*cc*_ = 0.01, *S*_*cv*_ = 0.1.

**Figure 11:**
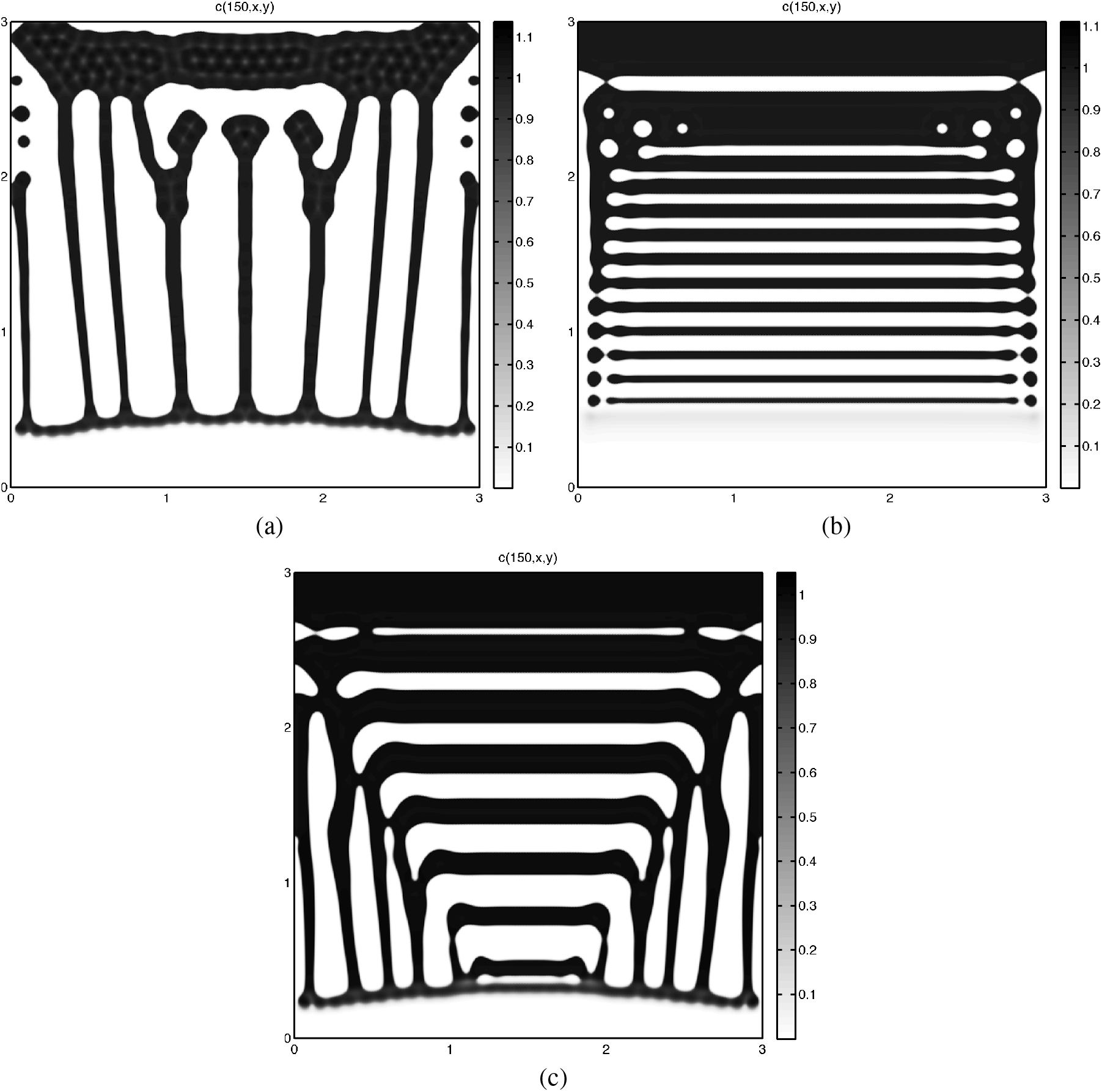
Plots showing the different patterns of cancer cell density *c*(*t, x, y*) using parameter set 𝒜 and *D*_*c*_ = 3.5*·*10^*−*4^ and *D*_*m*_ = 3.5*·*10^*−*4^, and varying the chemotactic coefficient and cell adhesion parameters. (a) “vertical stripe” pattern with *χ*_*m*_ = 0.04, *S*_*cv*_ = 0.1, *S*_*cc*_ = 0.1; (b) “horizontal stripe” pattern, with *χ*_*m*_ = 0.04, *S*_*cv*_ = 0.01, *S*_*cc*_ = 0.01; (c) mixed “horizontal and vertical stripe” pattern, with *χ*_*m*_ = 0.0305, *S*_*cv*_ = 0.1, *S*_*cc*_ = 0.01. All were taken at dimensionless time *t* = 150.

The relationship between vertical/horizontal patterns and high/slow invasion speeds, respectively, has been observed and studied by Köpf et al. Köpf et al. [2012]. Their work, using the Cahn-Hilliard equation to model the transfer of surfactant monolayers, demonstrated that low speeds produced striped patterns perpendicular to the direction of movement, whereas higher speeds resulted in striped patterns parallel to the direction of movement. We defer a detailed analysis of the pattern formation observed in our computational simulations to future work.

### 3.6 Model Extensions

We also outline potential further extensions of our current model. Other invasion studies, such as those by Gaggioli et al. Gaggioli et al. [2007] involving organotypic assays populated with stromal fibroblast cells, indicate that carcinoma cells migrate and invade by following these fibroblasts. The stromal fibroblasts act as leading cells, creating and leaving behind tracks in the extracellular matrix that facilitate the invasion of carcinoma cells as collective chains, maintaining epithelial characteristics. The structure of our mathematical model allows for straightforward extensions to simulate these other generic invasion systems, for instance, by explicitly incorporating the role of fibroblasts into the model, as inspired by the experiments of Gaggioli et al. Gaggioli et al. [2007]. In such an extension, fibroblasts (*f*) would be assumed to undergo random motility, proliferation, and production by external sources. Furthermore, we can account for different matrix components (*v*_*i*_, e.g., various collagens, fibronectin, vitronectin, laminin) and different matrix proteolytic enzymes (*m*_*j*_) that act on these specific matrix components. The extended system of equations is presented in (10).

Future development of this modeling approach (an *in silico* organotypic assay) should focus on estimating the model parameters from a single experimental setup and quantifying our adhesion parameters, specifically *S*_*cc*_ and *S*_*cv*_. Achieving these goals would enable the development of a robust quantitative modeling approach highly beneficial for experimentalists. Finally, we note that the model could also be adapted to examine and predict *in vivo* invasive cancer spread (which occurs on a different timescale) by modifying the adhesion parameters *S*_*cc*_ and *S*_*cv*_ to be time-dependent. Preliminary results in this direction have been reported by Domschke et al. Domschke et al. [2014], whose computational simulation results show qualitative similarities to the invasive growth patterns observed in various cancer types, such as tumor infiltrative growth patterns (INF) Ito et al. [2012], Masuda et al. [2012].

## 4 Conclusions

In this paper, we have presented an *in silico* model for cancer cell invasion based on experiments with *in vitro* organotypic assays, integrating the roles of proteolysis, cell adhesion, and cell migration. The *in silico* model was developed from a relatively simple mathematical model of cancer invasion with three key time- and space-dependent variables: cancer cell density, matrix density, and proteolytic enzyme concentration. This mathematical model comprises a system of partial differential equations with a nonlocal term governing adhesive interactions, comprehensively accounting for both cell-cell and cell-matrix adhesion. The model demonstrates biological realism, and its computational simulation results qualitatively capture *in vitro* experimental observations of local cancer cell invasion of tissue, where malignant tumor cells must detach from the primary tumor, degrade surrounding tissue, and migrate through it Liotta [1986].

Our *in silico* computational simulation results, presented in the **Results and Discussion** section, effectively demonstrate the influence of cell adhesion within the system. Varying cell adhesion parameters in the system of equations (9) yields diverse patterns of cell movement and invasion, and also affects the speed of movement. Experimentally, it has been proposed that, alongside the cancer cell environment, key properties of cancer cells, such as cell adhesion, can significantly influence the patterns of spread and growth Fidler [1978]. Utilizing our model with the nonlocal adhesive term, we can obtain *in silico* patterns that qualitatively replicate the invasive behavior observed in *in vitro* organotypic assays, such as the experiments by Martins et al. Martins et al. [2009] shown in Figs. 2, 3, and 8, by adjusting cell-cell and cell-matrix adhesion parameters. This adjustment allows for a degree of freedom in matching the simulated invasion speed or depth to experimental observations.

Our model produces results that are qualitatively comparable to those observed in several organotypic assay experiments. We are confident that our model can offer a deeper understanding of the processes involved in cancer cell invasion, particularly the complex but crucial adhesion processes that are vital for cancer studies and improving cancer treatment strategies.

